# Conditioned orienting predicts preference for, and affective response to, amphetamine after multiple exposures in male rats

**DOI:** 10.1101/2025.09.06.674656

**Authors:** Emily N. Hilz, Elena Morales-Grahl, Aditi Patel, Taylor Knight, Marie-H. Monfils, Hongjoo J. Lee

## Abstract

Conditioned orienting (OR) is a form of cue-directed behavior that is thought to reflect a behavioral phenotype of increased incentive-motivational processing of conditioned stimuli. Previous research has shown that OR also represents a cognitive phenotype that includes impaired attentional function and response inhibition, increased novelty seeking and, in females, resistance to extinction of conditioned place preference (CPP) for the stimulant drug amphetamine (AMP). The following experiments aim to increase our understanding of the OR cognitive phenotype in drug reward processing in male rats. In the first experiment, preference for AMP in males with or without an OR phenotype is assessed using CPP. In the second experiment, our understanding of OR as a predictor of AMP reward response is expanded by measuring potential behavioral mechanisms including affective and motor response to multiple AMP administrations. The results of the two experiments suggest that while the OR phenotype is not a predictor of resistance to extinction of AMP preference in males, OR as a discrete response variable predicts AMP preference after repeated exposures (i.e., AMP challenge). Moreover, we observed that rats with an OR phenotype had a higher positive affective response, measured via ultrasonic vocalizations (USVs) in the 50 kHz range, after multiple AMP administrations. OR scores predicted USVs but did not alter the gross motor response to AMP. Taken together, these results suggest that orienting behavior is a predictor of AMP preference in males, particularly after multiple exposures, and that one behavioral mechanism that could explain this is that male Orienters have a strong positive affective response to AMP.

## Introduction

Conditioned orienting is a form of cue-directed behavior that reliably occurs in a subset of animals that undergo Pavlovian appetitive conditioning. Over the course of repeated conditioned and unconditioned stimulus (CS-US) pairings, a subset of learners will acquire conditioned orienting responses (ORs), wherein the animal physically orients to the CS additionally – but not preferentially – to the US (Emily N. Hilz et al., 2019; E. N. Hilz et al., 2019; Hilz et al., 2023, 2022, 2021; Lee et al., 2011; Olshavsky et al., 2013a, 2013b). This phenomenon, first observed by Holland (Holland, 1977), is thought to represent increased incentive-motivational processing of CS information, such that the CS comes to acquire reinforcing properties of its own. This behavioral phenotype is similar in spirit to behavioral sign-tracking, which is also a type of cue-directed behavior that is thought to represent increased incentive-motivational processing of CS information (Flagel et al., 2009, 2007; Pitchers et al., 2015; Robinson and Flagel, 2009; Saunders and Robinson, 2011).

A substantial body of evidence links cue-directed behavioral phenotypes with increased preference for stimulant drugs like cocaine or amphetamine (Meyer et al., 2012; Tunstall and Kearns, 2015), higher consumption rates of these drugs (Beckmann et al., 2011), resistance to extinction of drug preference and/or consummatory behavior (Ahrens et al., 2016; E. N. Hilz et al., 2019; Morrison et al., 2015), and increased likelihood of relapse under various conditions (Everett et al., 2020; Saunders and Robinson, 2010; Versaggi et al., 2016). It is possible that these animals also receive more pleasure from stimulatory substances (Meyer et al., 2012; Tripi et al., 2017), measured in rats via ultrasonic vocalizations (USVs) within the 50 kHz range (Ahrens et al., 2013; Burgdorf et al., 2001). The theory that cue-directed behavior predicts increased likelihood of developing substance use disorder is strengthened by the relationship between cue-directed phenotypes with increased impulsivity and reduced attentional function (Hilz et al., 2021; Lovic et al., 2011; Olshavsky et al., 2014), which similarly predict a compulsive relationship with stimulant drugs (Diergaarde et al., 2008; Wit, 2009; Yates et al., 2012). We have previously shown that conditioned orienting as a discrete form of cue-directed behavior predicts increased impulsivity and higher distractibility in male rats (Olshavsky et al., 2014), and reduced cognitive flexibility and resistance to extinction of amphetamine conditioned place preference (CPP) in female rats (E. N. Hilz et al., 2019; Hilz et al., 2021). Here we seek to expand that characterization to determine if conditioned orienting similarly increases aspects of amphetamine preference in male rats, and further examine if such a relationship could be explained in part by increased pleasure experienced by male Orienters from amphetamine.

To do this, we conducted two experiments wherein adult male rats underwent Pavlovian food-light appetitive conditioning to determine the orienting phenotype. In the first experiment, the rats subsequently underwent amphetamine CPP followed by three days of place preference testing (with no amphetamine administration, so this also acted as extinction training) and then one day of amphetamine challenge. In the second experiment, the rats subsequently received two saline and amphetamine administrations over the course of four days and were monitored for locomotor behavior and USV response to the substances. These data help us determine if conditioned orienting represents a cognitive phenotype of increased vulnerability to substance abuse in male rats as it has been previously shown to in female rats, and provide new insight into a potential neuro-behavioral mechanism that may partially explain this vulnerability.

## Methods

### Subjects

A total of 62 male Sprague-Dawley rats (Envigo), aged ∼60+/- 5 days upon arrival, were used for the two experiments. The rats had a prior experience of undergoing an objection recognition test for a different study (Nemchek et al., 2021). All rats were housed in pairs on a 14:10 hour reverse light-dark cycle with lights off at 10AM. Water was always available *ad libitum*. Prior to Pavlovian conditioning rats were food-restricted for one week to reach ∼90% body weight and maintained this restriction throughout that procedure. After light-food conditioning, rats were allowed one week for weight recovery and were subsequently fed *ad libitum*. All procedures were conducted approximately 1-hour after the onset of the dark cycle under red light, were approved by the Institutional Animal Care and Use Committee at the University of Texas at Austin and were conducted in accordance with NIH guidelines.

### Apparati

Light-food conditioning occurred in a conditioning chamber measuring 30.5cm W / 25.5cm L / 30.5cm H (Coulbourn Instruments, Whitehall, PA). Briefly, the chambers were constructed of clear acrylic front and back walls, steel-rod floors, and aluminum sides and ceiling. Food-pellets (45mg TestDiet, Richmond, IN) were dispensed into a foodcup on the right wall from an external magazine, and entries into the foodcup were measured via breaks in an infrared beam at the opening. A 2-watt bulb was located 20cm above the foodcup, and illumination of this light served as the CS for conditioning. Chambers were enclosed in sound- and light-attenuating boxes (Coulbourn Instruments, Whitehall, PA). Mounted inside the boxes but outside the chambers were digital video cameras (KT&C USA, Fairfield, NJ) used to monitor orienting activity during conditioning.

Conditioned place preference procedures occurred in a conditioning chamber similar in construction but of different dimensions (50.8cm W / 25.5cm L / 29.2cm H; Coulbourn). Walls had no perforations and an aluminum wall with a retractable door bisected the chamber into separate compartments. The door was closed during amphetamine and saline conditioning, and open during baseline / preference testing / reinstatement. Black and white paper was secured to the acrylic walls of each compartment such that one side (left) was “dark” and the other (right) was “light”. A 2-watt red light located 22.5cm above the floor provided ambient light in the dark compartment. A down-facing camera was mounted atop the dark compartment which recorded time spent by the rat within each compartment. Subtracting time spent in the dark chamber from the total session length allowed determination of time spent in the white chamber and subsequent generation of place preference scores. Boxes were enclosed in sound- and light-attenuating boxes.

AMP-Saline USV responses were measured in 43.5cm L / 21.5cm W / 20cm H standard polystyrene housing cages with filter tops and no bedding. USVs were recorded with CM16 ultrasonic microphones and AvisoftRECORDER (Avisoft Bioacoustics, Berlin, Germany), and digitized at a 200 kHz sampling rate with 16-bit resolution using UltraSoundGate system (Avisoft Bioacoustics). The cages were placed atop a cart in open air under red light; microphones were placed outside the cage facing a small open port approximately 10cm high on the short edge of the box, and video cameras were mounted outside the cage facing the long edge. A line was drawn halfway along the long edge to bisect the cage into two 21.5cm sections and used to count cage crossing. Two rats underwent the procedure at a time, in separate cages approximately 5 feet apart.

### Procedures

#### Experiment 1: Conditioned orienting as a predictor of amphetamine CPP

##### Pavlovian Conditioning

The first cohort of male rats (*n* = 38) underwent Pavlovian conditioning procedures where a 10-second light CS was paired with delivery of one food pellet US. On the first day, rats were trained to retrieve the food pellet from the magazine where 30 pellets were delivered on a 60-second fixed inter-trial interval (ITI) over a 30-minute session. The following day, rats were habituated to the light CS. This consisted of 8 CS-only presentations where the light was illuminated without delivery of the food US, followed by 8 CS-US pairings where the CS preceded delivery of the US. The habituation session lasted ∼35- minutes and trials occurred on a variable ITI of 120s +/- 60s. The rest of the conditioning sessions occurred over the subsequent 3 days, and each consisted of 16 CS-US presentations on the same variable ITI schedule.

The Pavlovian conditioning procedure was used to determine the conditioned orienting behavioral phenotype for the rats. Orienting responses (ORs) were measured via video recording over the habituation and conditioning sessions and later quantified by an independent observer over a 15-second sampling period partitioned into 3 distinct intervals: 5-seconds prior to the CS illumination (preCS), first 5-seconds of the CS illumination (CS1), and second 5- seconds of the CS illumination (CS2). ORs were assessed every 1.25-seconds, allowing for up to 4 ORs for each sampling period. ORs were operationally defined as a rearing response towards the CS where the front legs of the rat lifted from the floor, excluding grooming behavior or reaching into the food magazine. Because the CS fully illuminated the conditioning chamber, orientation of the rat during ORs was not considered. ORs typically occur in CS1 and food cup approach occurs predominantly in CS2 (E. N. Hilz et al., 2019; Lee et al., 2011; Olshavsky et al., 2014)) – baseline preCS scores were subtracted from CS1 and CS2 behavior for orienting and foodcup behavior, respectively.

### Conditioned place preference

CPP procedures similar to those in Hilz et al., 2019a were used. CPP consisted of four phases: baseline, amphetamine (AMP) and saline conditioning, preference testing / extinction, and challenge. Baseline was a 15-minute session wherein rats were placed into the conditioning chamber on the dark side and allowed to roam freely between the dark and light sides through the open retractable door; baseline determined unconditioned place preference scores. The subsequent conditioning procedures were biased such that amphetamine was paired with the less preferred context. Conditioning sessions consisted of an intraperitoneal (i.p.) injection of either d-amphetamine (AMP; 2.0 mg/kg, Sigma-Aldrich) or 0.9% sterile saline prior to placement within the conditioning chamber with the retractable door closed. The first conditioning session paired AMP with confinement to the less-preferred context for 30 minutes. The second paired saline with confinement to the more-preferred context for 30 minutes. This procedure was repeated such that rats received two AMP and two saline conditioning sessions over 4 days. Two days after the final conditioning session, preference testing occurred over three consecutive days and, as no AMP i.p. injection occurred, acted additionally as extinction training. Preference / extinction tests were 15-minute sessions wherein rats were placed into the conditioning chamber on the dark side and allowed to roam freely between the dark and light sides through the open retractable door. Finally, an AMP challenge was given to assess AMP CPP after the extinction training sessions. This occurred two days after preference / extinction testing. Rats were given AMP i.p. injection prior to placement within the dark chamber and allowed to roam freely through the open retractable door for 15 minutes.

### Experiment 2: Conditioned orienting and hedonic response to amphetamine

#### Pavlovian Conditioning

The second cohort of rats (*n* = 24) underwent Pavlovian conditioning procedures identical to those described in Experiment 1 to again determine OR phenotype and conditioned approach behavior.

#### AMP Treatment and Ultrasonic Vocalizations

The goal of the second experiment was to determine if OR phenotype predicts changes in AMP-sensitivity measured by ultrasonic vocalizations (USVs) and locomotor response (i.e., cage crossing and rearing). To do this, rats underwent a 5-day AMP-Saline treatment schedule where 24-hours passed between each session. The first session was habituation, where rats were placed for 15-minutes into a clean cage and allowed to habituate to the room and cage. The next session was the first saline treatment (Sal1); rats were i.p. injected with 0.9% sterile saline and video and USVs were recorded. Recordings took place for 30 minutes after AMP or saline treatment; the subsequent sessions were Amp1, where rats were i.p. injected with 2.0 mg/kg of d-amphetamine, Sal2 (i.e., second saline treatment), and finally Amp2 (i.e., second AMP treatment). Two rats were run concurrently for each session.

### Track USF

Ultrasonic vocalizations were analyzed using TrackUSF, an open-source software that breaks up ultrasonic vocalizations into 6-ms fragments (USFs) and then groups these USFs based on their frequency using t-distributed Stochastic Neighbor Embedding (t-SNE) analysis (Netser et al., 2022). The threshold was set to 2.7 based on Netser et al. (2022), and automatic clustering was performed using TrackUSF’s automatic clustering algorithm. 5 distinct USF clusters were identified (Fig. 1A), and USF clusters with > 50% overlap in frequency were combined for analysis; a random selection of representative spectrograms from each cluster were visually assessed by an independent observer and classified as either USVs or noise (Fig. 1B). Generally, noise can be identified by its structure: USVs are short spectrogram bursts that begin and end within a certain frequency range (e.g., in the affective range, USVs typically occur in 50 +/- 20 kHz range; Knutson et al., 2002; Willey et al., 2009), while noise typically shows a “peak”-like structure beginning at 0 kHz. Of the 5 clusters, clusters 1 and 4 were identified as noise based on their structure and removed from further analysis.

**Fig. 1.**
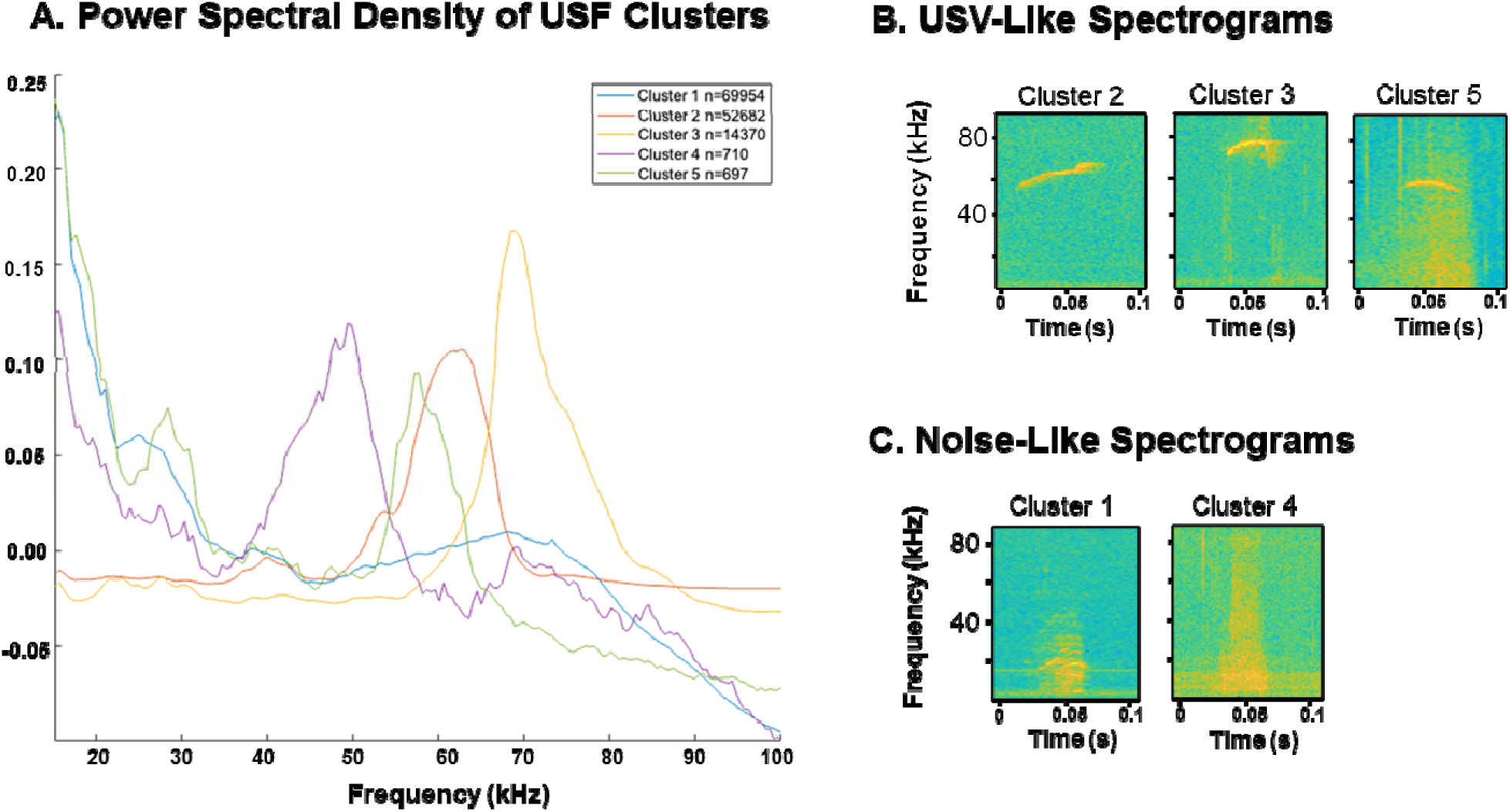
**A.** Mean Power Spectral Density (PSD) of Ultrasonic Vocalization Fragment (USF) clusters’ frequency. The normalized power of each cluster is shown on the y-axis, and the frequency (Hz x 10^4^) at which that power is concentrated is displayed on the x-axis. N = number of USFs in each cluster. **B**. Representative spectrograms from clusters 2, 3, and 5 with USV-like structures. Clusters 2 and 5 overlapped significantly in frequency and were combined for analysis, while cluster 3 was analyzed separately. **C.** Representative spectrograms from clusters 1 and 4 with noise-like structures. Clusters 1 and 4 were excluded from analyses.

Clusters 2 and 5 were determined to have > 50% overlap in frequency, occurred in the 50 +/- 20 kHz range, and were combined into a single outcome score for analysis. Cluster 3 was comprised of USFs that occurred between 60 and 80 kHz. Because it had little overlap with clusters 2 and 5, and a comparatively high frequency (with approximately 50% of the USFs above 70 kHz), cluster 3 was analyzed separately.

### Statistical Analyses

All analyses were conducted in R 4.3.1 using the base stats package unless otherwise stated (R Core Team, 2023).

### Hierarchal clustering to determine OR phenotype

Orienters and Nonorienters were categorized based on hierarchal clustering on the mean OR level at the end of Pavlovian conditioning (i.e., last 8 trials) for Experiment 1 and Experiment 2, separately. First, a Euclidean distance matrix was computed on the mean OR scores using the ‘dist()’ function; hierarchal clustering by Ward’s method (i.e., “ward.D2”) was performed on this matrix using the ‘hclust()’ function. The Within-Cluster Sum of Squares was calculated and scree-plotted to determine the optimal number of clusters using ‘ggplot()’ to determine the elbow point. This value was used to categorize the rats binarily into Orienters (i.e., cluster of high OR scores) and Nonorienters (i.e., cluster of low OR scores).

### Behavioral analyses

The Pavlovian conditioning of foodcup and orienting behavior, CPP for amphetamine, and USV and locomotor response to amphetamine were analyzed using within-subjects 2x2 ANOVAs for repeated measures data or between-subjects 2x2 ANOVAs for non-repeated measures data. Normality and homogeneity of variance in the data were first confirmed using Shapiro-Wilk test of normality and Bartlett’s test of homogeneity of variance. Data was compared using 2x2 ANOVA with the common factor “Phenotype” (i.e., Orienter or Nonorienter). For Pavlovian conditioning, the factor “Block” indicated acquisition of responses over time (i.e., blocks of 8 trials, 4 days). For CPP, the factor “Day” indicated preference for amphetamine over the extinction/testing days (i.e., days 1-3) and the amphetamine reinstatement challenge (i.e., day 4). For the USV and locomotor response, the factor “Treatment” indicated the injection schedule of either saline (i.e., SAL1 or SAL2) or amphetamine (i.e., AMP 1 or AMP2). Significant effects were assessed using appropriate *post hoc* analyses: Bonferroni correction for repeated measures data, otherwise Tukey HSD; partial eta squared (n_p_^2^) is provided as a measure of effect size for significant comparisons with *post hoc* analyses. For n_p_^2^, 0.01 is considered a small effect size, 0.06 is considered a medium effect size, and 0.14 is considered a large effect size (Richardson, 2011).

## Results

### Acquisition of conditioned responses

Data from both experiments were combined for the analysis of Pavlovian conditioning (*n* = 69). The first block of 8 trials in the Pavlovian conditioning paradigm represents unconditioned foodcup and orienting responses from habituation where the light CS was presented without the food pellet US. Analyses were therefore restricted to block 2-8. Over the conditioning blocks, foodcup behavior was higher in CS2 compared to CS1, indicated by a significant main effect of CS (*F*(1,793) = 4.73, *p* = 0.03) and Block (*F*(6,793) = 11.29, *p* < 0.001), as well as a significant interaction between CS x Block (*F*(6,793) = 22.72, *p* < 0.001). Fig. 2A shows that FC entries were higher by the end of conditioning in CS2 compared to CS1 (*F*(1,122) = 48.53, *p* < 0.001, n_p_^2^ = 0.28; Tukey HSD: *p* < 0.001). Inversely, ORs were higher in CS1 compared to CS2, indicated by a significant main effect of CS (*F*(1,793) = 14.75, *p* = 0.001), Block (*F*(6,793) = 4.68, *p* = 0.001), and interaction between CS x Block (*F*(6,793) = 12.20, *p* < 0.001). Fig. 2B shows that ORs were higher by the end of conditioning in CS1 compared to CS2 (*F*(1,122) = 8.91, *p* < 0.001, n_p_^2^ = 0.19; Tukey HSD: *p* < 0.001).

**Fig 2.**
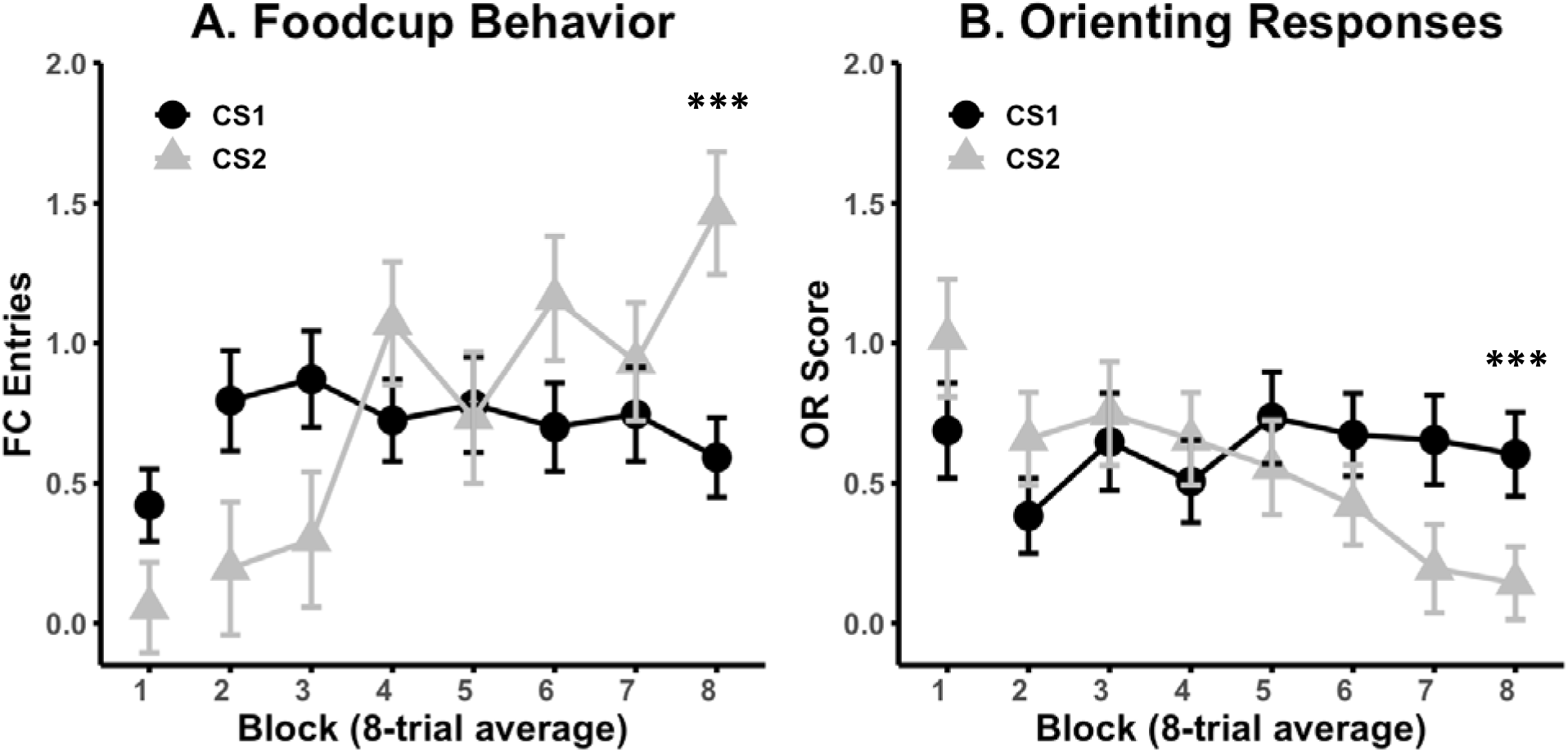
Acquisition of conditioned responses between CS1 and CS2. Numbers on the x-axis represent blocks of 8- trials; the first block was unpaired light conditioned stimulus (CS) presentations. **A.** Mean entries into the FC receptacle +/- SEM over the course of food-light Pavlovian conditioning for both experimental cohorts (data combined; *n* = 62). Male rats had higher FC behavior during the second portion of the CS presentation, CS2, and higher FC response in CS2 compared to CS1 at the end of conditioning (i.e., block 8). **B.** Mean orienting response (OR) score +/- SEM over the course of food-light Pavlovian conditioning for both experimental cohorts (data combined). Male rats had higher OR behavior during the first portion of the CS presentation, CS1, and higher OR scores in CS1 compared to CS2 by the end of conditioning (i.e., block 8). *** = *p* < 0.001

### Determination of orienting phenotype

For experimental cohort 1 (*n* = 38), hierarchal clustering resulted in a dendrogram with 4 clusters of varying *n* sizes (Fig. 3A). An elbow point was identified at 2 clusters (Fig. 3B), suggesting that the data was best represented binarily. Rats in cluster 1, where the average OR level was 1.13, were identified as “Orienters”; rats in cluster 2, where the average OR level was 0.13, were identified as “Nonorienters”. For experimental cohort 2 (*n* = 24), hierarchal clustering also resulted in a dendrogram with 4 clusters of varying *n* sizes (Fig. 3C). An elbow point was identified at 2 clusters (Fig. 3D), suggesting that the data was best represented binarily. Rats in cluster 1, where the average OR score was 0.009, were identified as “Nonorienters”; rats in cluster 2, where the average OR score was 1.125, were identified as “Orienters”.

**Fig. 3.**
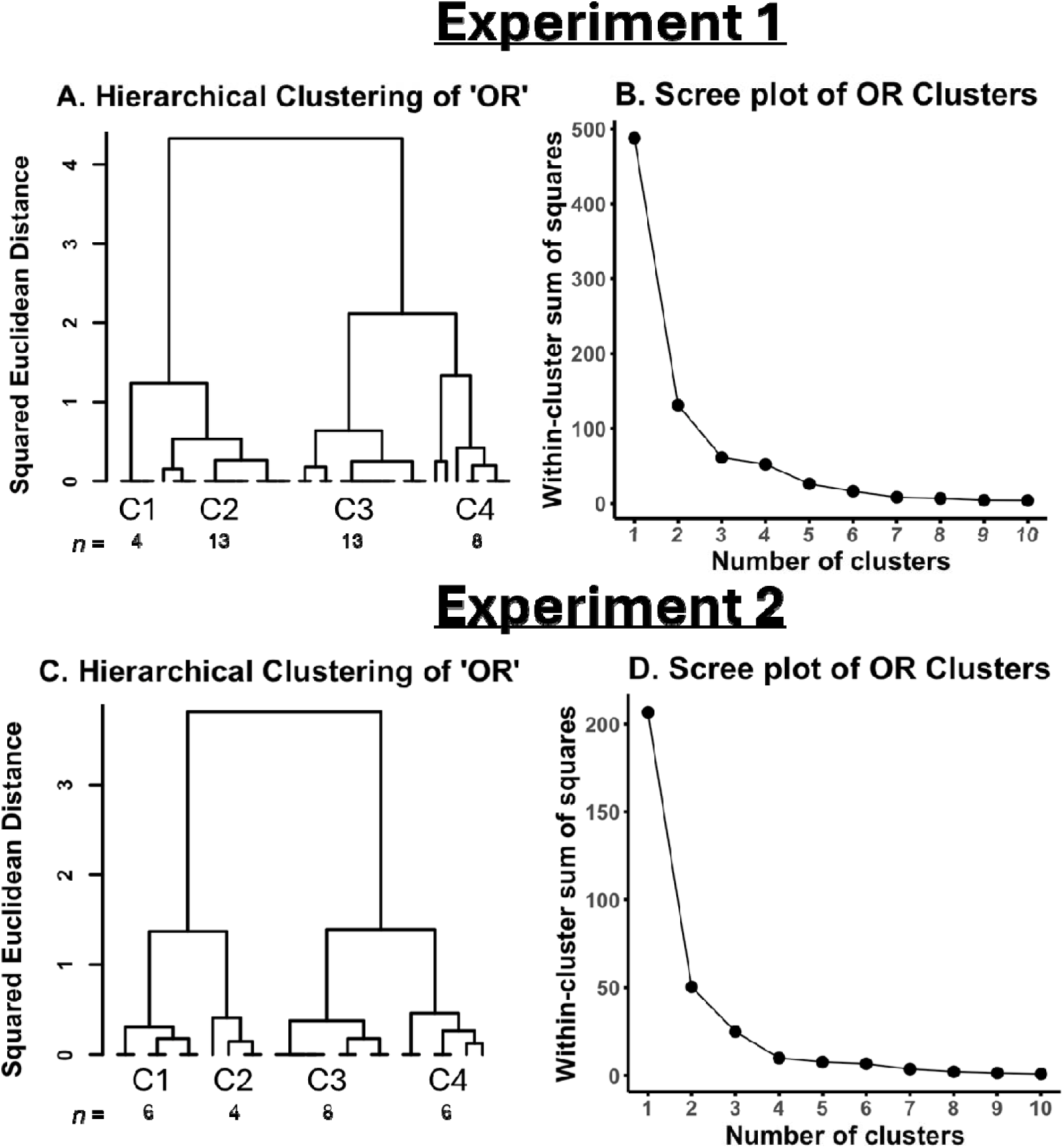
Determination of conditioned orienting phenotype. **A.** Dendrogram showing unique clusters of orientin responses (ORs) determined by the squared Euclidean distance between datapoints for the first experimental cohort. Four unique clusters were identified (C 1 – 4), and the number of individuals in each cluster is given (n). **B.** Scre plot showing the within-cluster sum of squares (WCSS) for 10 potential sample clusters (y-axis). A distinct elbow occurs at point 2, indicating that segregating the data into two clusters is the optimal way to minimize cluster number and the variance within each cluster. **C.** Dendrogram showing unique clusters of orienting responses (ORs) determined by the squared Euclidean distance between datapoints for the second experimental cohort. Four uniqu clusters were identified (C 1 – 4), and the number of individuals in each cluster is given (n). **D.** Scree plot showing the WCSS for 10 potential sample clusters (y-axis). A distinct elbow occurs at point 2, indicating that segregating the data into two clusters is the optimal way to minimize cluster number and the variance within each cluster.

Data from both experiments were combined for the analysis of Pavlovian conditioning by orienting phenotype (*n* = 69). We have previously found that rats acquire FC behavior similarly over Pavlovian conditioning regardless of OR phenotype; however, in this data we observed a significant effect of OR Phenotype on acquisition of FC behavior indicated by a significant interaction of Phenotype and Block (*F*(6,360) = 2.70, *p* = 0.01) although no main effect of Phenotype on its own (*F*(1,60) = 1.41) was detected. When analyzing the average FC scores at block 8, FC behavior was lower among Orienters compared to Nonorienters (*F*(1,60) = 6.01, *p* = 0.01, n_p_^2^ = 0.09, Tukey HSD: *p* = 0.02; Fig. 4A). A similar delineation, although in the opposite direction, was observed for acquisition of ORs. A significant main effect of Phenotype (*F*(1,60) = 21.69, *p* < 0.001) and a significant interaction between Phenotype and Block (*F*(6,360) = 3.67, *p* = 0.001) indicated differential acquisition of ORs by Phenotype, and when analyzing the average OR level at block 8, OR level was higher in Orienters compared to Nonorienters (*F*(1,60) = 49.35, n_p_^2^ = 0.45, *p* < 0.001, Tukey HSD: *p* < 0.001; Fig. 4B).

**Fig. 4.**
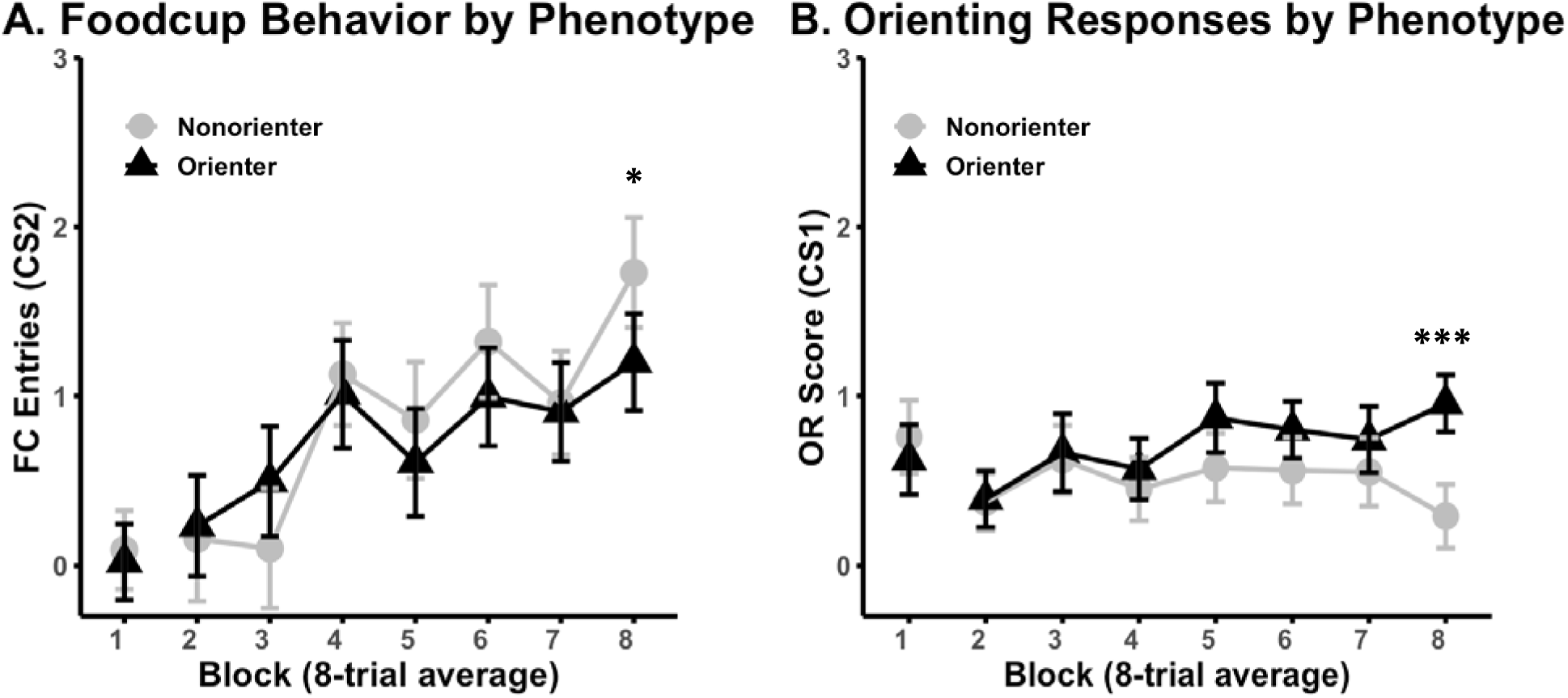
Comparing acquisition of conditioned responses between Orienters (*n* = 27) and Nonorienters (*n* = 35). Numbers on the x-axis represent blocks of 8-trials; the first block was unpaired light conditioned stimulus (CS) presentations. **A.** Mean entries into the FC receptacle +/- SEM over the course of food-light Pavlovian conditioning for both experimental cohorts (data combined). Male Orienters had a significantly lower FC response by the end of conditioning (i.e., block 8). **B.** Mean orienting response (OR) score +/- SEM over the course of food-light Pavlovian conditioning for both experimental cohorts (data combined). Male Orienters had a significantly higher OR score by the end of conditioning (i.e., block 8). * = *p* = 0.02, *** = *p* < 0.001

### Experiment 1 - Amphetamine conditioned place preference

The first experimental cohort went on to undergo AMP CPP using a biased procedure. We observed that the side, light or dark, within which rats underwent AMP CPP was an important factor in overall AMP preference. For the first three days of testing / extinction training, a significant main effect of side was detected (*F*(1,34) = 9.29, n_p_^2^ = 0.21, *p* = 0.004) and a non-significant main effect of Day (i.e., test/extinction days 1-3; *F*(2,68) = 2.94, n ^2^ = 0.08, *p* = 0.06). When we considered AMP preference by side in separate ANOVAs, we found that rats conditioned to AMP in the light side (*n* = 22) showed a significant reduction in AMP place preference over testing/extinction days indicated by a main effect of Day (*F*(2,40) = 3.92, n_p_^2^ = 0.16, *p* = 0.02), while those conditioned to the dark (*n* = 16) side did not (*F*(2,28) = 0.36). For the light side, *post hoc* comparisons revealed that AMP preference was significantly lower at EXT 3 compared to EXT 1 (*p* = 0.02; Fig. 5A); however, there were no main nor interaction effects of Phenotype during the test/extinction days for rats conditioned to either the light or dark side (*p* > 0.1 for all comparisons). At AMP challenge, AMP place preference did not differ based on Phenotype (*F*(1,34) = 2.10) nor side (*F*(1,34) = 0.007; Fig. 5B).

**Fig. 5.**
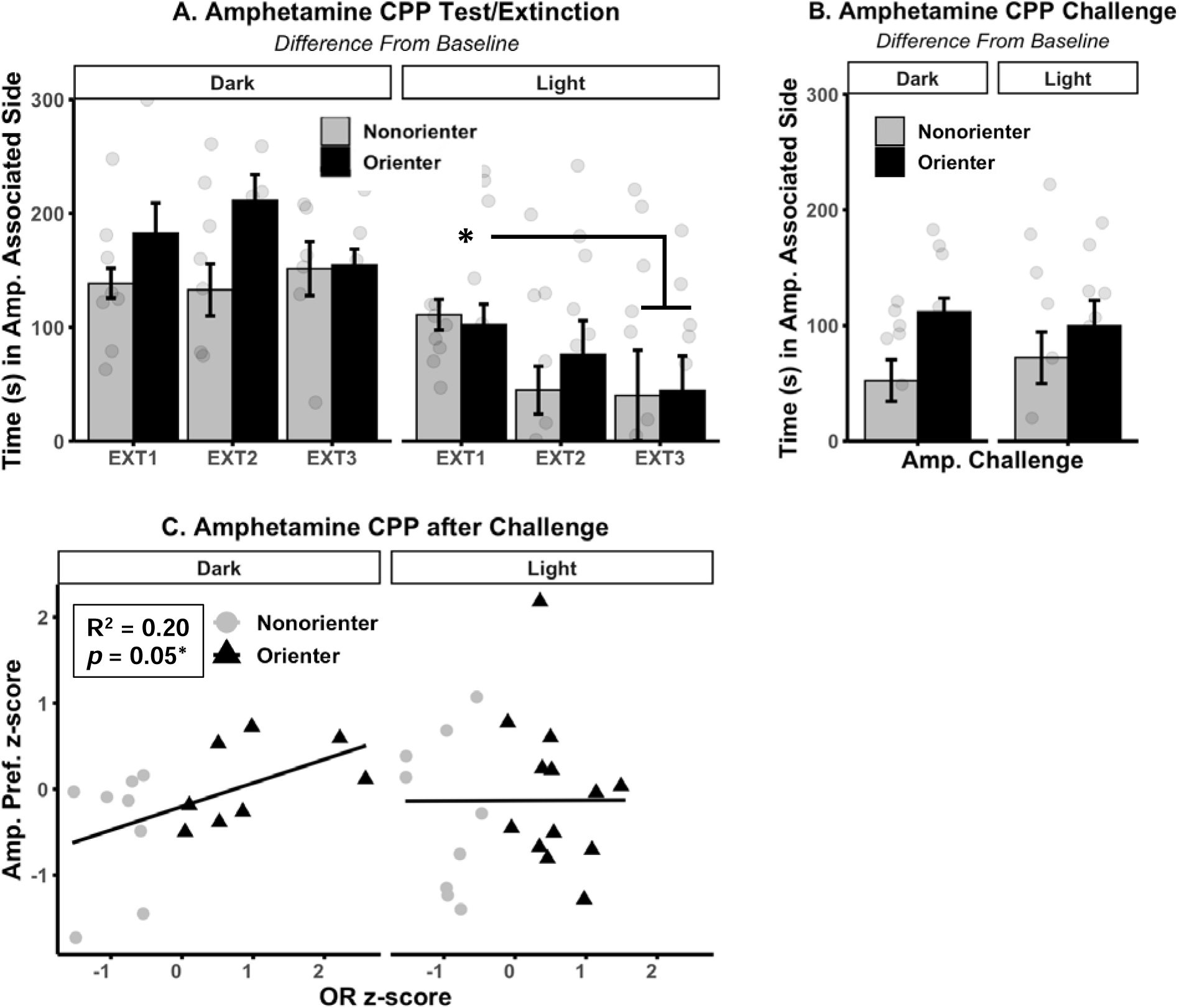
Experiment 1 - Amphetamine (Amp.) conditioned place preference (CPP). **A.** Mean time (s) as a difference from baseline spent in the Amp. associated context +/- SEM over the three testing/extinction (EXT 1 – 3) days. The data is partitioned into rats conditioned to Amp. in the dark (*n* = 16) or light context (*n* = 22); Amp. preference was significantly higher in the dark-conditioned rats. There was no effect of the conditioned orienting (OR) phenotype on Amp. preference over testing/extinction. **B.** Mean time (s) as a difference from baseline spent in the Amp. associated context +/- SEM after Amp. challenge. There was no effect of the OR phenotype on Amp. Preference (Pref.) after challenge. **C.** Linear regression showing relationship between OR z-score and Amp. preference z-score after challenge. OR score positively predicted Amp. preference after challenge in rats conditioned to the dark context.

Although there were no effects oberseved when considering the OR Phenotype as a factor, we did observe a linear relationship between OR scores at the end of Pavlovian conditioning (i.e., average of block 8) and place-preference for the AMP-associated context after the AMP challenge. Specifically, using a linear regression model with OR score as the independent variable and AMP place preference as the dependent variable for rats conditioned in the dark context, we found that OR scores positively predicted AMP preference after challenge (*F*(1,14) = 4.67, *p* = 0.05, R^2^ = 0.20; Fig. 5C). This was not the case for rats conditioned in the light context, among which no relationship was observed.

### Experiment 2 - USV response to amphetamine

The second experimental cohort underwent USV monitoring after multiple AMP or saline administrations. The sum of clusters 2 and 5 had over 50% overlap of frequency and were detected primarily in the 50 +/- 20 kHz range; from clusters 2 and 5, a significant main effect of Treatment (*F*(3,66) = 10.51, n_p_^2^ = 0.32, *p* < 0.001) indicated that the number of USV fragments were significantly higher after both AMP treatments compared to saline (*p* < 0.002 all). A main effect of Phenotype (*F*(1,22) = 3.66, n_p_^2^ = 0.14, *p* = 0.07) and the interaction between Phenotype and Treatment (*F*(3,66) = 2.32, n_p_^2^ = 0.10, *p* = 0.08) did not reach conventional levels of significance; however, exploratory *post hoc* comparisons on the interaction in the omnibus ANOVA suggested that Orienters may have emitted more USVs than Nonorienters after the second AMP exposure (*p* = 0.05). We confirmed this by conducting an ANOVA on only the “Amp2” condition and found a signficant main effect of Phenotype (*F*(1,22) = 8.17, *p* = 0.009, n_p_^2^ = 0.27, Tukey HSD: *p* = 0.009; Fig. 6A). We also found a significant linear relationship between OR scores at the end of Pavlovian conditioning and the number of USV fragments detected in the 50 kHz range; using a linear regression model with OR score as the independent variable and USV fragments from the combined 2 and 5 cluster as the dependent variable, we found that as OR score increased, so did the number of USVs emitted after the second AMP administration (*F*(1,22) = 11.41, *p* = 0.003, R^2^ = 0.31; Fig. 6B).

**Fig. 6.**
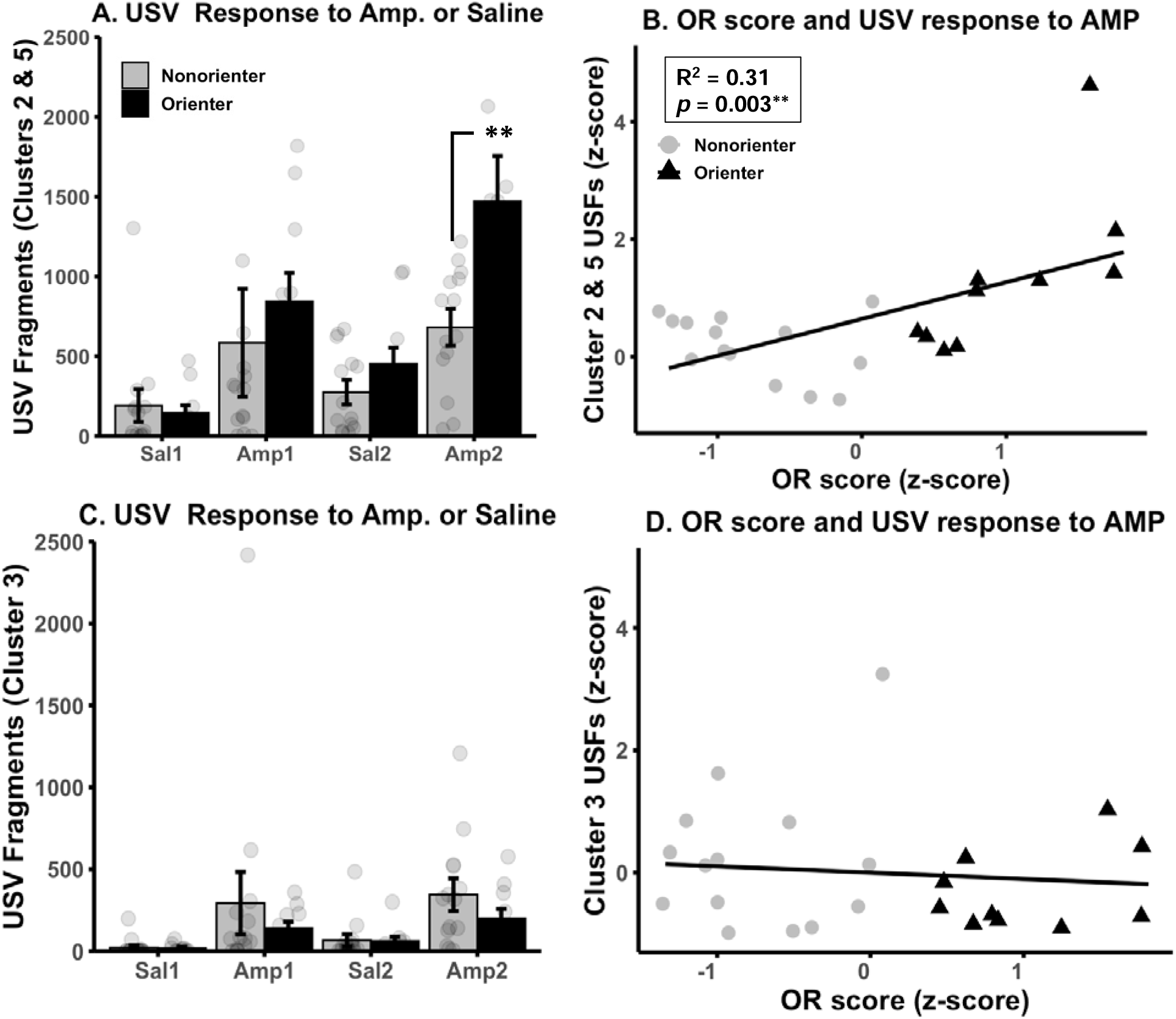
Experiment 2 – Ultrasonic Vocalizations (USVs) after amphetamine or saline. **A.** Mean number of USV fragments summed from clusters 2 and 5 or TrackUSF +/- SEM over the two amphetamine and saline administrations (i.e., Sal 1 & 2, Amp 1 & 2). Rats emitted more USVs at Amp 1 and 2 compared to Sal 1 and 2; additionally, Orienters (*n* = 10) emitted more USVs at Amp 2 compared to Nonorienters (*n* = 14). **B.** Linear regression showing relationship between OR z-score and USV fragments z-score after at Amp 2. OR score positively predicted the number of USV fragments detected by TrackUSF after the second amphetamine administration. ** = *p* = 0.009

Cluster 3 represented USFs in the 60 – 80 kHz range and overlapped < 50% with clusters 2 and 5. From cluster 3, a significant main effect of Treatment (F(3,66) = 5.96, n_p_^2^ = 0.21, p = 0.001; Fig. 6C) indicated that the number of USV fragments were significantly higher after both AMP treatments compared to saline (*p* = 0.05 and *p* = 0.04, chronologically). There were no main effects of Phenotype (*p* > 0.1) nor interactioin between Phenotype and Treatment (*p* > 0.1), nor was a linear relationship detected between cluster 3 USFs and OR score (*p* > 0.1; Fig. 6D).

We also considered cage crossing and rearing as locomotor responses to AMP or saline administration, and found a significant main effect of Treatment on both cage crossing (*F*(3,66) = 48.89, *p* < 0.001; Fig. 7A) and rearing (*F*(3,66) = 62.94, *p* < 0.001) such that both response measures were higher after AMP administration than saline (all *post hoc* comparisons *p* < 0.001). However, only rearing was affected by the OR Phenotype. A significant interaction was observed between Phenotype and Treatment (*F*(3,66) = 3.90, n_p_^2^ = 0.15, *p* = 0.01) such that that rearing was lower among Orienters compared to Nonorienters at Amp2 (*p* = 0.01; Fig. 7B). This may reflect a marginal but non-significant increase in rearing among Nonorienters between Amp1 and Amp2 (*p* = 0.08).

**Fig. 7.**
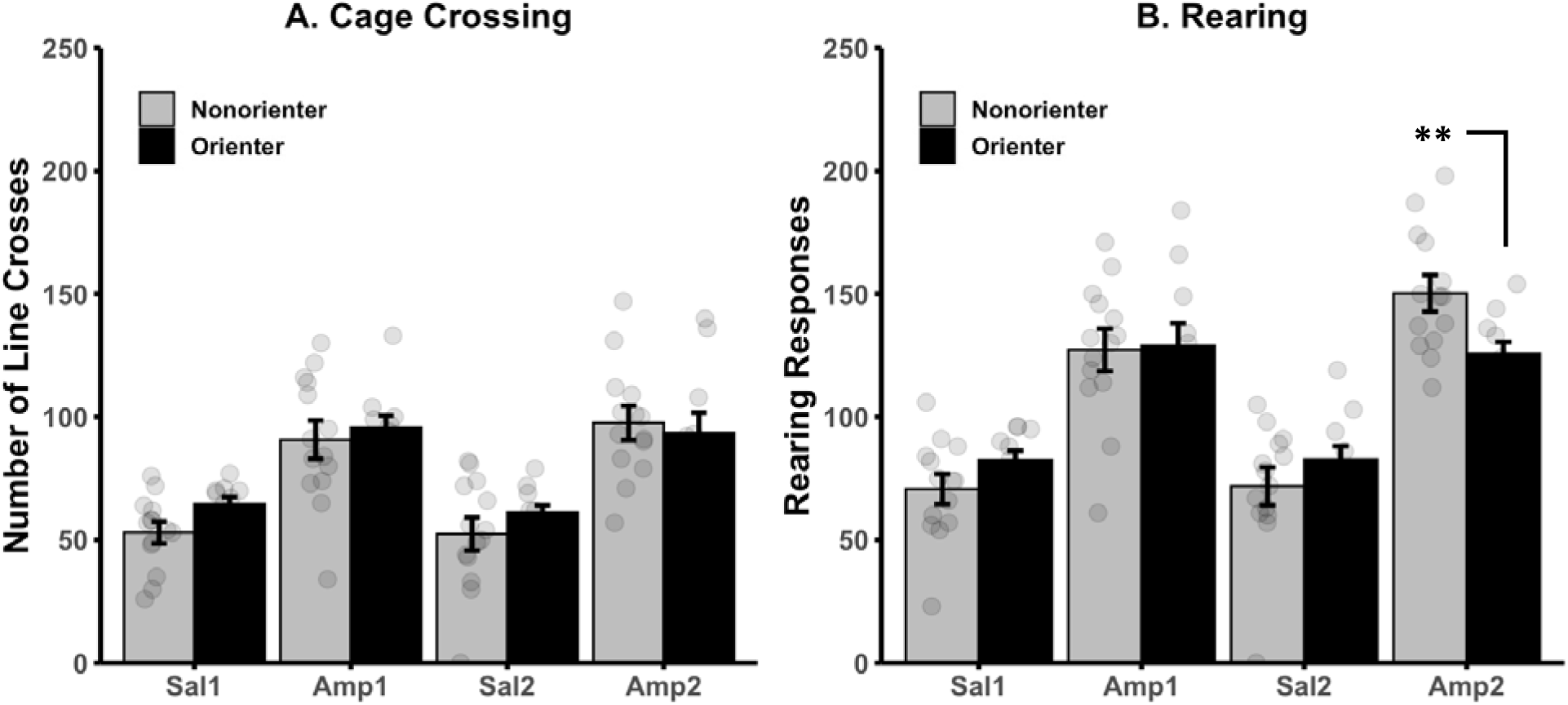
Experiment 2 – Motor responses after amphetamine or saline. **A.** Mean number of cage (line) crossings +/- SEM over the two amphetamine and saline administrations (i.e., Sal 1 & 2, Amp 1 & 2). Rats crossed the cage more frequently at Amp 1 and 2 compared to Sal 1 and 2; there was no effect of the orienting phenotype. **B.** Mean number of rearing responses +/- SEM over the two amphetamine and saline administrations. Rats reared more frequently at Amp 1 and 2 compared to Sal 1 and 2; additionally, Orienters (*n* = 10) reared less after the second amphetamine administration compared to Nonorienters (*n* = 14). ** = *p* = 0.01

## Discussion

The data presented provides evidence that conditioned orienting in male rats may predict increased preference for the stimulant drug amphetamine (AMP), and may also increase the rewarding and/or pleasureable effects of AMP administration. In experiment 1, we conditioned rats to AMP in either a light or dark context based on their baseline place preference. In females, we have previously observed resistance to extinction of AMP place preference among Orienters compared to Nonorienters (E. N. Hilz et al., 2019). Unlike females, we saw no effect of the orienting phenotype on AMP place preference over the course of the testing/extinction days.

### AMP preference in Orienters after extinction and challenge

There is a notable difference between this experiment and the one conducted in females. In the female experiment, all rats had a higher baseline preference for the dark context and so were conditioned to AMP only in the light context. In this experiment, a higher proportion of male rats had a baseline preference for the light context, and were conditioned to AMP in either the light or dark context based on that baseline preference. The rats conditioned to AMP in the dark context show no observable extinction curve; however, the rats conditioned to AMP in the light context did extinguish their AMP preference over multiple days. Cue-directed behavioral phenotypes have been shown to predict resistance to exinction of reward-seeking behaviors in our lab and others (Ahrens et al., 2016; Fitzpatrick et al., 2019; E. N. Hilz et al., 2019; Morrison et al., 2015). Here we observe that while male Orienters exhibited resistance to extinction of AMP preference in the dark context, the same was true for Nonorienters. When AMP was paired with the light context both Orienters and Nonorienters extinguished their AMP preference behavior. Because AMP preference is significantly higher among rats conditioned in the dark context, it is possible and likely that the aversive qualitities of the light stimulus (Arrant et al., 2013) contribute to extinction of AMP preference, and perhaps male Orienters are sensitive to this aversive quality in a way that differs from female Orienters.

While we observed no group effect of the orienting phenotpe on AMP preference after challenge, we did observe a relationship between the individual levels of orienting response (OR) and AMP preference. As OR score increased, so did preference for the AMP-associated context after AMP challenge. This indicated that while male Orienters may not be resistant to extinction of AMP preference, they may be more resposive to the stimulatory or appetitive effects of direct AMP administration particularly after multiple administrations.

### Orienting increases affective USV response to AMP

The evidence that male orienters are more sensitive to AMP after repeated adminstrations is strengthened by our second experiment in which we administered AMP or saline to male rats two times and measured ultrasonic vocalizations (USVs) and locomotor behavior in response. USVs in rats are considered reflective of affect or emotional state: USVs in the 50 +/- 20 kHz range represent positive affect, and have been used as a proxy measurement of ‘liking’ (e.g., have a positive affective response to) amphetamine, among other rewarding substances and stimuli (Ahrens et al., 2013, 2009; Burgdorf et al., 2007, 2001; de Oliveira Guaita et al., 2018). We used a novel software, TrackUSF, to automatically detect and classify USVs based on frequency (Netser et al., 2022). The software segregates recorded sounds into 6 millisecond fragments and clusters the fragments based on freuqency; from there, an independent observer must examine the fragments within clusters to determine if the cluster has collected USV-type sounds, or noise. We cannot say that our USV fragment data represents the total number of USVs emitted by the rats nor can we identify the structure of the calls for more nuanced analysis. However, we were able to quantify the gross effects of AMP administration on the total number of sounds produced in the 50 kHz range by rats that have been exposed to AMP or saline.

The patterns of these sounds are in-line with what others have found when manually measuring USVs after AMP: the number of USVs were significantly higher in rats administered AMP compared to saline (Ahrens et al., 2013, 2009; Burgdorf et al., 2001; Mulvihill and Brudzynski, 2019). This was true for both USV fragement clusters analyzed, although the effect was more robust in clusters 2 and 5. Additionally, we observed an effect of the OR phenotype on the number of cluster 2 and 5 USV fragments emitted after AMP such that male Orienters emitted more USVs after the second AMP administration compared to Nonorienters, and OR score was a positive predictor of the USV response to the second AMP administration. This may indicate that male Orienters may have an increased hedonic response to amphetamine – more ‘liking’ – than do Nonorienters, which has been similarly observed in male sign-trackers after cocaine administration (Meyer et al., 2012; Tripi et al., 2017). Notably, these effects were not oberseved in the analysis of cluster 3 USV fragements. Cluster 3 was comprised to high frequency (70 +/- 10 kHz) sounds that had a USV-like structure; although 50% fell above the range of what we typically consider the “affective” range of USVs, we cannot say this cluster of USFs represents a unique response class based on the data. USVs are detected up to 80 kHz in experiments that measure reward and pleasurable stimulation (Shimoju et al., 2020; Simola et al., 2018), although comparatively high frequency USVs are also measured in social interaction tests in rats (A. Berz et al., 2022; A. C. Berz et al., 2022; Willey et al., 2009). Because our experimental procedure was conducted on two rats concurrently in separate cages 5 feet apart, it is possible that cluster 3 represents a social interaction-type USV. Without direct measurement, which was beyond the goals of the experiment, the precise function of cluster 3 USFs is conjecture on our parts. What is clear is that these higher frequency USV fragments were not affected by OR phenotype, and less were emitted over the course of testing.

### Orienting does not affect AMP-induced locomotor behavior

We did not find consistent effects of the OR phenotype on locomotor response, suggesting effects of the OR phenotype are restricted to the affective qualities of AMP. We observed that AMP administration increased these locomotor responses, as expected; however, modualtion of locomotor response by OR phenotype was differential between the horizontal and vertical planes. Specifically, cage crossing was unaffected, but Nonorienters engaged in more rearing after the second AMP exposure than did Orienters. In sign- versus goal-tracking experiments, cocaine adminstration increases locomotor activity in goal-trackers at comparatively high doses (Flagel et al., 2008). Others have observed no differences in locomotor response to stimulant drugs based on cue-directed behavior, novelty-seeking, nor in rats selectively bred to produce high or low USV response to AMP (Garcia and Cain, 2016; Mu et al., 2009; Tripi et al., 2017). Our data suggests that the orienting may similarly predict a positive neurobehavioral response to AMP that is delineated from drug-induced locomotion.

In conclusion, these experiments provide preliminary evidence that conditioned orienting, as a discrete form of cue-directed behavior, predicts preference for and affective response to amphetamine uniquely in male rats. By extending previous results from female to male rats and observing differential behavioral responses between the sexes, we underscore the necessity of considering sex differences in drug-related behaviors. This offers a more comprehensive understanding of factors that may contibute to compulsive drug use and drug-seeking behaviors, as well as contributing to our understanding of the neurobiological phenomena that underscore this. Although not measured here, we and others have shown repeatedly that integrity of the midbrain dopamine system is necessary for cue-directed behavior to occur and mediates the attribution of incentive salience to cues (Bonansco et al., 2018; Flagel et al., 2007; Lee et al., 2011, 2006; Pitchers et al., 2017; Singer et al., 2016); future research would benefit from investigating how male and female Orienters recruit dopamine after exposure to drugs of abuse to determine if sex differences exist in this neurobiological substrate that may explain the unique but completmentary effects of the OR phenotype between the sexes.

